# NuGraph: Graph-Based Reasoning over 3D Primitives for Nucleus Segmentation Correction

**DOI:** 10.64898/2026.05.16.725603

**Authors:** Mingzhi Wang, Peng Liu, Yi Zhao, Bingzhang Wang, Jia Wan, Liqiang Nie, Donglai Wei

## Abstract

Correcting segmentation errors in large-scale 3D nuclei reconstructions requires reasoning about which fragments belong to the same nucleus across densely packed regions. Existing correction methods rely on local pairwise fragment matching, which cannot resolve the global topology of nuclear clusters and fails to recover missing morphology. We propose NuGraph, a graph-based reasoning framework that operates over atomic 3D primitives obtained by decomposing erroneous masks. NuGraph encodes primitive geometry via a 3D point-cloud backbone and performs global relational reasoning through graph attention, capturing inter-primitive dependencies across entire clusters rather than isolated pairs. A primitive–proposal contrastive loss aligns local primitive features with nucleus-level semantics, improving grouping accuracy in dense regions. The resulting proposals are then refined by a shape-refinement network that predicts signed distance fields to restore smooth morphology. To train without manual error annotations, we develop a self-supervised data engine that synthesizes realistic segmentation errors from clean nuclei labels. To benchmark correction at brain scale, we curate NucEMFix, the first brain-wide EM benchmark of nuclei error cases across FAFB and MICrONS (8,000+ annotated error nuclei). NuGraph attains 87.99% F1 on NucEMFix-F (FAFB) and 86.20% on NucEMFix-M (MICrONS), outperforming both re-segmentation baselines (e.g., +8.6% over nnU-Net) and pairwise correction methods, while reducing curation effort by over 100× relative to manual proofreading. Code and data are available at https://mingzhiwang618.github.io/NucEMFix.

## I. Introduction

Accurate 3D nuclei segmentation provides the foundation for a wide range of biological studies—from development and regeneration to large-scale anatomy [1], [2]. In neuroscience, nuclei serve as spatial anchors linking cellular identity to circuit reconstruction and cross-scale integration [3]–[6], and segmentation quality directly impacts downstream analyses [7]–[9]. As shown in Fig. 1, landmark EM datasets such as FAFB and MICrONS have unlocked cellular-scale mapping at unprecedented scope, yet their released nuclei segmentations still contain errors despite extensive proofreading [10], [11].

**Fig. 1.**
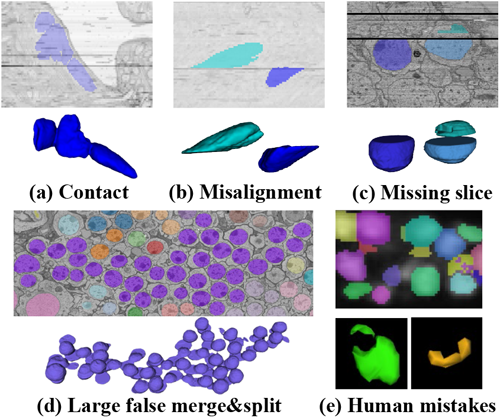
Long-tail nuclei segmentation errors in public 3D microscopy datasets: (a) ambiguous contact between nuclei yielding false merges; (b) misalignment with discontinuous objects; (c) missing slices with axial gaps; (d) merge–split hybrids from misalignment and dense packing; (e) annotation artifacts.

The *last mile* of 3D nuclei segmentation is dominated by rare, heterogeneous failures—false merges in densely packed regions due to weak boundary cues [12] and false splits from acquisition artifacts disrupting z-axis continuity [13]. Correcting these errors demands *reasoning* about which fragments belong to the same nucleus, yet most prior correction methods rely on local pairwise fragment matching [14]–[16]. These approaches face a fundamental limitation: local pairwise decisions cannot resolve the global topology of densely packed nuclear clusters. Unlike neurons, where skeletons provide explicit merge candidates, nuclei lack such structural priors, making local reasoning insufficient for complex merge–split cases. Furthermore, pairwise matching cannot recover missing morphology caused by axial gaps or EM artifacts, which is critical for downstream cell typing and morphological analysis.

To address this, we propose NuGraph, a graph-based reasoning framework that operates over atomic 3D primitives decomposed from erroneous masks, reducing diverse error types to a common structural representation. Graph attention reasons across all primitives in a cluster, capturing the global topological context that local pairwise methods miss, while a primitive–proposal contrastive loss aligns local primitive features with nucleus-level semantics. A shape-refinement head then predicts signed distance fields to restore smooth morphology. To eliminate manual error annotation, we develop a self-supervised data engine that synthesizes realistic segmentation errors from clean labels.

To benchmark correction at brain scale, we curate **NucEMFix**, a systematic benchmark of nuclei error cases across FAFB (Drosophila) and MICrONS (Mouse), encompassing over 8,000 annotated error nuclei. Across both datasets, NuGraph consistently outperforms re-segmentation baselines and pairwise correction methods.

### The main contributions are

- **Graph-based nucleus reasoning**. We formulate nucleus error correction as relational reasoning over atomic 3D primitives, using graph attention with primitive–proposal contrastive alignment to overcome the limitations of local pairwise methods.
- **NucEMFix benchmark**. The first brain-wide EM benchmark (FAFB, MICrONS) for nuclei correction, featuring 8,000+ annotated error nuclei with diverse failure modes and a self-supervised data engine for scalable training.
- **State-of-the-art performance**. NuGraph achieves the highest F1 scores on NucEMFix, outperforming resegmentation and pairwise correction baselines.

### II. Related work

#### A. 3D Nuclei Instance Segmentation

Voxel-wise approaches such as 3D U-Net [17] and nnU-Net [18] learn semantic masks but struggle to separate tightly packed instances. Shape-aware methods introduce geometric priors: StarDist [19] parametrizes nuclei as star-convex polyhedra, while Cellpose [20] predicts spatial gradient flows. Despite these advances, deploying general-purpose segmenters on large-scale, anisotropic EM volumes remains challenging due to the interplay of low axial resolution, image artifacts, and dense cellular packing, producing characteristic false merges and false splits that require dedicated correction.

Public benchmarks span modalities and species, including *C. elegans* [21], rodent and human tissue [1], [22], and cancer cell lines [23]. NucMM [24] provides large-scale microCT and EM nuclei volumes, and NIS3D [25] offers densely annotated 3D fluorescence data. These resources focus on segmenting all nuclei from scratch. Our **NucEMFix** complements them by targeting the last-mile error space in brain-scale EM, supplying curated error cases for correction benchmarking.

#### B. Segmentation Error Correction in Connectomics

Automated proofreading has primarily targeted *neuron* reconstruction via multiscale error detection [26], topological priors [14], [27], or skeleton-based tracing [28]–[30]. Pairwise approaches classify fragment pairs for merging using point-cloud features [15] or learned edge networks [16]. Graph-based formulations such as multicut and agglomeration [31], [32] optimize global partitioning with learned edge weights but typically initialize edges only between adjacent supervoxels, limiting their ability to reason over densely packed structures. Transferring these paradigms to nuclei faces fundamental challenges. Nuclei are volumetric blobs rather than curvilinear structures, so simply linking fragments cannot recover complete morphology. Nuclei also form dense clusters where local adjacency is ambiguous and global context is essential. NuGraph addresses both: graph attention captures global topology across entire clusters, and SDF refinement restores smooth morphology in postprocessing.

## III. Methodology

### A. Overview

We formulate nucleus error correction as a relational reasoning problem over atomic 3D primitives. As illustrated in Fig. 2, NuGraph operates in three stages: (1) *shape decomposition*, which decomposes erroneous masks into atomic primitives 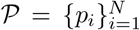 (2) *graph-based relational reasoning*, which groups primitives into nucleus proposals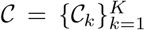 global graph attention; and (3) *shape refinement*, which restores complete morphology by predicting signed distance fields for each proposal, yielding the final output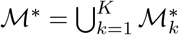.

**Fig. 2.**
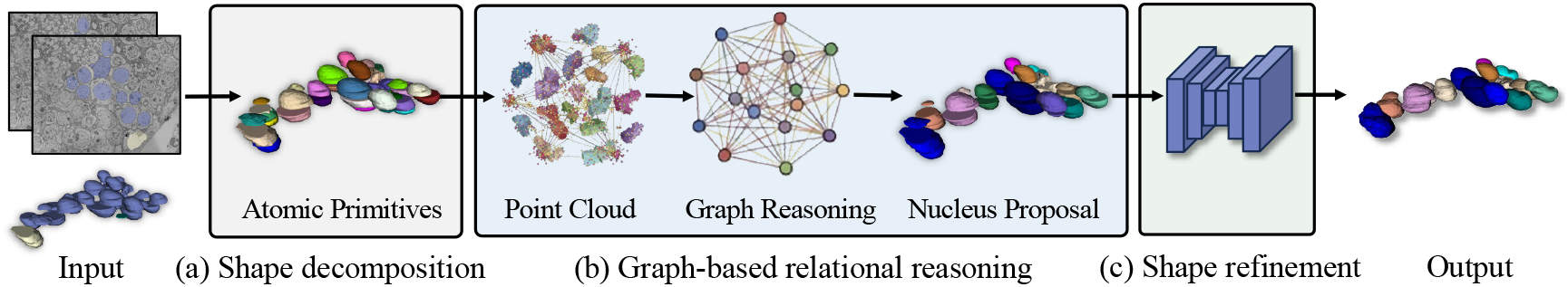
Overview of the NuGraph framework. (a) **Shape decomposition**: erroneous masks are split into atomic 3D primitives via EDT-based watershed. (b) **Graph-based relational reasoning**: point-cloud features of primitives are encoded and reasoned over by a graph network to group them into nucleus proposals. (c) **Shape refinement**: a 3D U-Net predicts signed distance fields to restore complete nucleus morphology. A self-supervised data engine synthesizes realistic errors from clean labels for training.

### B. Shape Decomposition

To reduce diverse error types to a common representation, we decompose erroneous masks into atomic 3D primitives. Given a mask ℳ^0^, we apply realignment preprocessing [33], remove severely degraded slices, compute the Euclidean Distance Transform (EDT), and apply watershed on the negated EDT to split at geometric bottlenecks. Specifically, local maxima of the EDT are identified as watershed seeds using a minimum peak distance of *d*_min_ = 10 voxels, ensuring that only sufficiently separated geometric centers generate distinct primitives. This produces an over-complete set of primitives 𝒫= {*p*_*i*_}. Small fragments below volume *v*_min_ are merged with neighbors, reframing error correction as a structured grouping problem.

### C. Graph-Based Relational Reasoning

As illustrated in Fig. 2(b), the core of NuGraph is relational reasoning over primitives: determining which fragments belong to the same nucleus by capturing global context across entire clusters, rather than making isolated pairwise comparisons. The detailed architecture of this stage is further illustrated in Fig. 3.

**Fig. 3.**
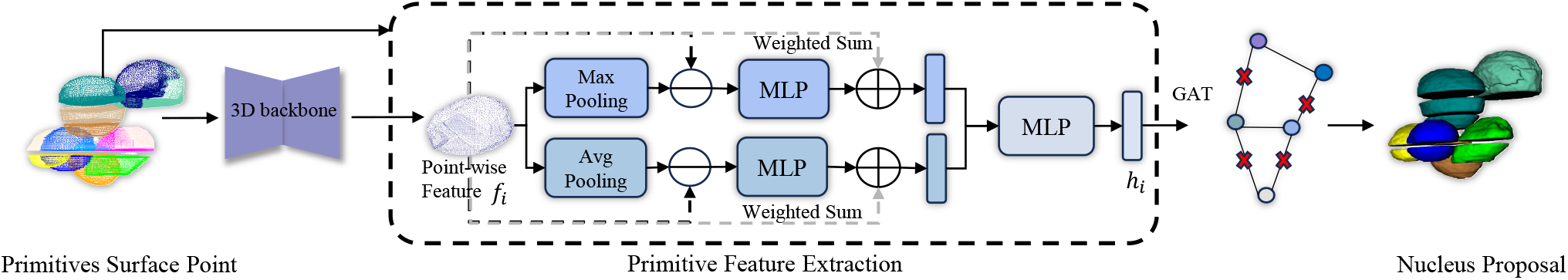
Graph-based relational reasoning: surface points of primitives are encoded by a 3D backbone, then aggregated into primitive-level features via dual-path attention pooling. Graph attention networks capture global topological context, with primitive–proposal contrastive learning aligning local primitives with global proposal structure. Edge classification groups primitives into nucleus proposals.

#### Geometric Feature Encoding

Given primitives𝒫= {*p*_*i*_}, we sample surface points from all primitives as a unified input and process them through a point-cloud encoder to obtain point-wise features 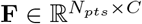:

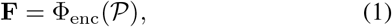

where Φ_enc_ denotes the 3D Sparse-UNet [34] backbone, and the input surface point coordinates are normalized before encoding.

#### Primitive Feature Extraction

To aggregate point-level features 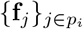into a compact descriptor **h**_*i*_ for each primitive *p*_*i*_, we employ a dual-path residual attention pooling module to capture both local salient and overall contextual features.

Specifically, let **f**_*j*_ ∈ ℝ^*C*^ denote the point-wise feature of point *j* extracted from **F**, and 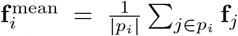 and 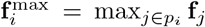the mean- and max-pooled features of primitive *p*_*i*_, respectively. We compute residual features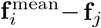 **f**_*j*_ and 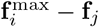, which quantify the relative contribution and spatial saliency of each point *j* within primitive *p*_*i*_.

Two separate MLPs process these residuals to generate attention weights, which are normalized via softmax. The final primitive descriptor is obtained by concatenating the attention-weighted aggregations from both paths and passing through a fusion MLP:

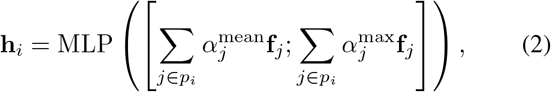

where 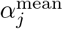 and 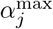are the softmax-normalized attention weights for the mean and max pooling paths, respectively.

#### Graph Reasoning

Given the primitive descriptors {**h**_*i*_} obtained from the previous module, we construct an undirected graph𝒢 =(𝒱,ℰ) where each node *v*_*i*_ ϵ 𝒱corresponds to a primitive *p*_*i*_ with feature **h**_*i*_. Two nodes are connected by an edge if the minimum pairwise distance between their surface points is below a threshold *d* (measured in voxels):

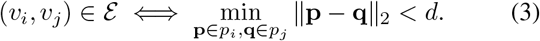

To capture high-order topological context, we employ an *L*-layer Graph Attention Network (GAT) [35] to iteratively refine node features. Specifically, given initial node features 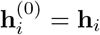, the *l*-th layer computes:

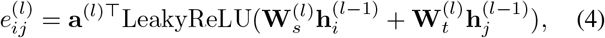

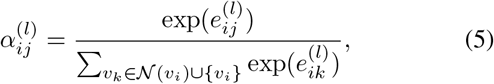

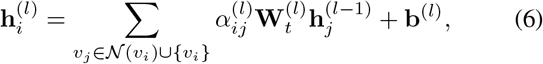

where 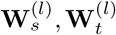, are learnable linear projection matrices for the source and target nodes, respectively, **a**^(*l*)^ is a learnable attention vector, **b**^(*l*)^ is a bias term, and 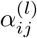 denotes the normalized attention coefficient from node *v*_*j*_ to node *v*_*i*_ at layer *l*. We then obtain the refined node features 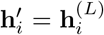, where we set *L* = 2 by default.

Given these refined features, we classify each edge to determine whether the connected primitives belong to the same nucleus. For each edge (*v*_*i*_, *v*_*j*_) ∈ ℰ, we concatenate the node features and pass them through an MLP classifier:

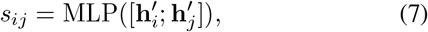

where *s*_*ij*_ is the predicted logit score. Edges with predicted probability *σ*(*s*_*ij*_) *> τ*_*e*_ are retained to form a pruned graph𝒢^′^ =(𝒱,ℰ^′^), where we set *τ*_*e*_ = 0.5 as threshold under our balanced positive/negative edge sampling:

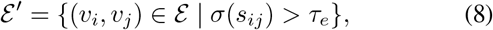

We then extract connected components from 𝒢^′^ as nucleus proposals 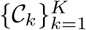, where each proposal 𝒞_*k*_ represents a set of primitives predicted to belong to the same nucleus instance.

#### Primitive–Proposal Contrastive Alignment

Graph reasoning refines primitive features via spatial interactions, but many primitives remain geometrically incomplete, forming irregular sheet-like structures due to missing slices. These fragments lack holistic nucleus semantics, limiting graph-based reasoning. As shown in Fig. 4, to bridge local primitive features with global nucleus-level semantics, we introduce a primitive– proposal contrastive loss. For each nucleus proposal 𝒞_*k*_, we aggregate features from all component primitives using the same attention pooling module to obtain a proposal embedding **c**_*k*_. The contrastive objective aligns each primitive feature with its corresponding proposal while enforcing separation between proposals:

**Fig. 4.**
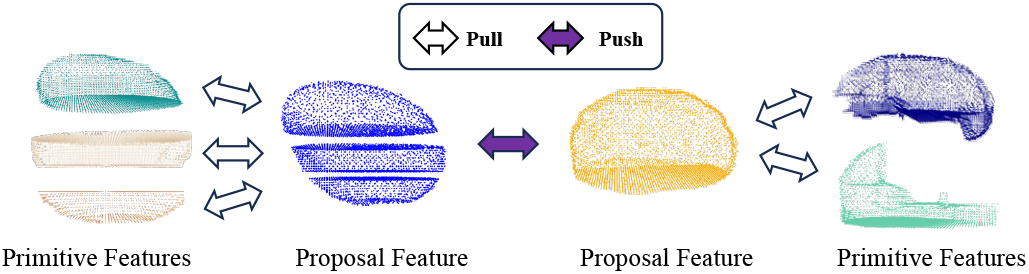
Primitive–proposal contrastive alignment. Each primitive is pulled toward its proposal embedding while proposal embeddings are pushed apart, bridging local primitive features with nucleus-level semantics.

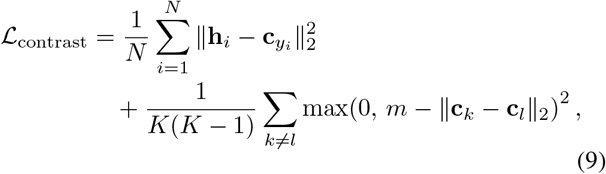

where *y*_*i*_ is the ground-truth nucleus index of primitive *i*, and *m* is the margin enforcing minimum separation between proposal embeddings. During training, proposals 𝒞_*k*_ are formed by grouping primitives according to ground-truth nucleus labels (rather than predicted edges) to provide a stable target for alignment; at inference they are produced by the predicted edge classifier as in Eq. (8).

#### Overall Training Objective

The complete loss combines edge classification with contrastive learning:

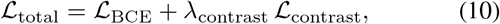

where *λ*_contrast_ is a weighting coefficient, and the edge classification loss ℒ_BCE_ is defined as

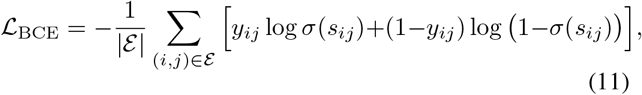

where *y*_*ij*_ ∈ {0, 1} denotes the ground truth label indicating whether primitives *i* and *j* belong to the same ground-truth nucleus, and *σ*(*·*) is the sigmoid function.

#### Self-Supervised Training

To learn relational reasoning without manual error annotations, we synthesize realistic segmentation errors from clean nuclei masks. We randomly place 1–20 nuclei in a shared 3D space, iteratively displacing each along random directions until it contacts existing nuclei, creating stochastic overlaps that mimic false merges. The nuclei are then truncated along the axial direction and fragmented via EDT-watershed—consistent with the input representation stage—to generate labeled primitives for self-supervised training.

### D. Shape Refinement

After relational reasoning produces nucleus proposals 𝒞_*k*_, a shape-refinement network restores complete morphology by predicting a continuous signed distance field (SDF). For each voxel *v*, the SDF is defined as *S*(*v*) = −*d*(*v, ∂*Ω) if *v*ϵ Ω and *S*(*v*) = *d*(*v, ∂*Ω) otherwise, where *v* ∈ Ω is the foreground volume and *d*(*·*) the Euclidean distance to the boundary. The SDF is normalized to [*−*1, 1].

As shown in Fig. 5, a 3D U-Net takes each proposal and its corresponding image patch as input and jointly predicts an SDF *Ŝ*_*k*_ and a foreground mask 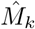. Training pairs are synthesized from clean labels via stochastic slice masking (removing adjacent mask slices) and patch masking (occluding foreground blocks). The training objective combines mask supervision, SDF regression, and total variation regularization for smooth surfaces:

**Fig. 5.**
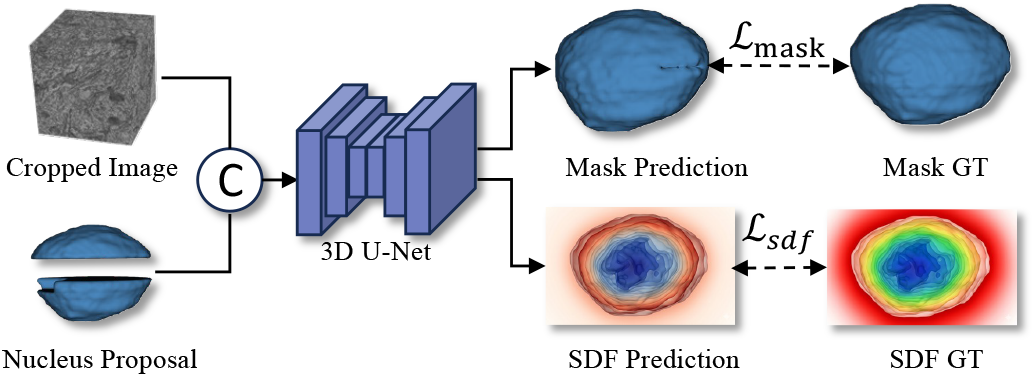
Shape refinement: a 3D U-Net predicts signed distance fields from each proposal and its image patch, restoring complete nucleus morphology. A self-supervised data engine synthesizes realistic error scenarios from clean nuclei labels for training.

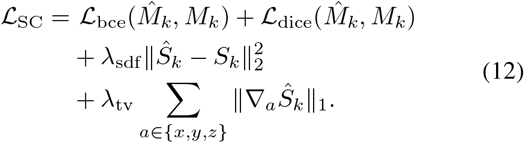

where *λ*_sdf_ and *λ*_tv_ are balancing weights.

## IV. NucEMFix Benchmark

### A. Dataset Source

We curate NucEMFix from two landmark public datasets to ensure diversity in resolution, species, and staining: the Full Adult Fly Brain (FAFB) [36] and the MICrONS dataset (1 mm^3^ mouse visual cortex) [4]. For FAFB, we use the released whole-brain nuclei segmentation [37]; for MICrONS, we sample from the central subvolume to mitigate boundary artifacts.

### B. Hierarchical Data Curation

Given the petabyte scale of these volumes (containing millions of nuclei), manual inspection of every instance is infeasible. We therefore devised a two-stage curation pipeline to systematically harvest high-value error cases.

#### Automated Candidate Mining

To efficiently locate errors within the PB-scale data, we implemented a heuristic screening protocol based on geometric priors:

- *Merge Candidates*: We targeted instances that exceeded biologically plausible volume or bounding box dimensions. These large candidates were subjected to multi-step morphological erosion; instances that fractured into multiple connected components were flagged as probable false merges (aggregates of multiple nuclei).
- *Split Candidates:* We identified potential false splits primarily via voxel count ranking. Instances falling significantly below the minimum expected volume of a complete nucleus were flagged as fragments. These candidates represent “shattered” parts of a nucleus or isolated slices resulting from z-axis discontinuities, often lacking their complementary halves in the initial segmentation.

#### Manual Inspection and Voxel-level Correction

Following automated mining, we conducted rigorous manual correction by three trained annotators (*∼*250 hours).

- *Brain-wide Systematic Review:* Beyond the automatically flagged candidates, annotators performed a systematic inspection of all segmented nuclei across the full extent of the datasets using Neuroglancer [38]. This exhaustive review enabled us to discover error cases that evaded the heuristic screening. Critically, annotators documented the specific failure mode for each error case, categorized by root cause (e.g., misalignment, boundary ambiguity). This fine-grained error taxonomy provides valuable insights into the failure modes of current segmentation methods.
- *Voxel-level Correction:* All identified error cases—both from automated mining and systematic review—were exported to ITK-SNAP [39] for voxel-level editing.

#### Expert Consensus Review

To ensure label consistency, two senior experts double-reviewed all corrected samples. Only cases with unanimous consensus entered the final benchmark.

### C. Dataset Statistics

Across both subsets, errors are long-tailed: a few very large merge cases dominate, while many small split fragments populate the tail (Fig. 6). NucEMFix-F also contains larger nuclei on average than NucEMFix-M, leading to more extensive error regions. As shown in Tab. I, NucEMFix-F consists of 3584 *×*8192 *×*14336 voxels at 80 *×*64 *×*64 nm^3^ resolution, containing 106, 978 neuronal nuclei, among which we annotated *>* 3,000 error cases affecting *>* 7,300 (*∼* 7%) nuclei. NucEMFix-M comprises 1632 *×*4096*×* 6144 voxels at the same resolution, with 77, 475 nuclei and 400 error cases affecting *>* 750 nuclei. Note that the counts above denote *nuclei affected*, since a single error case may involve multiple nuclei, especially in false merge cases. The mouse brain is substantially larger than that of Drosophila, resulting in more sparsely distributed nuclei and consequently fewer segmentation errors compared to FAFB. In MICrONS, we observed that most errors occur in the vicinity of blood vessels, affecting endothelial and perivascular cell nuclei. These nuclei often exhibit irregular morphologies, leading to false merges or fragmented instances, and thus represent a key target for future correction strategies.

**Fig. 6.**
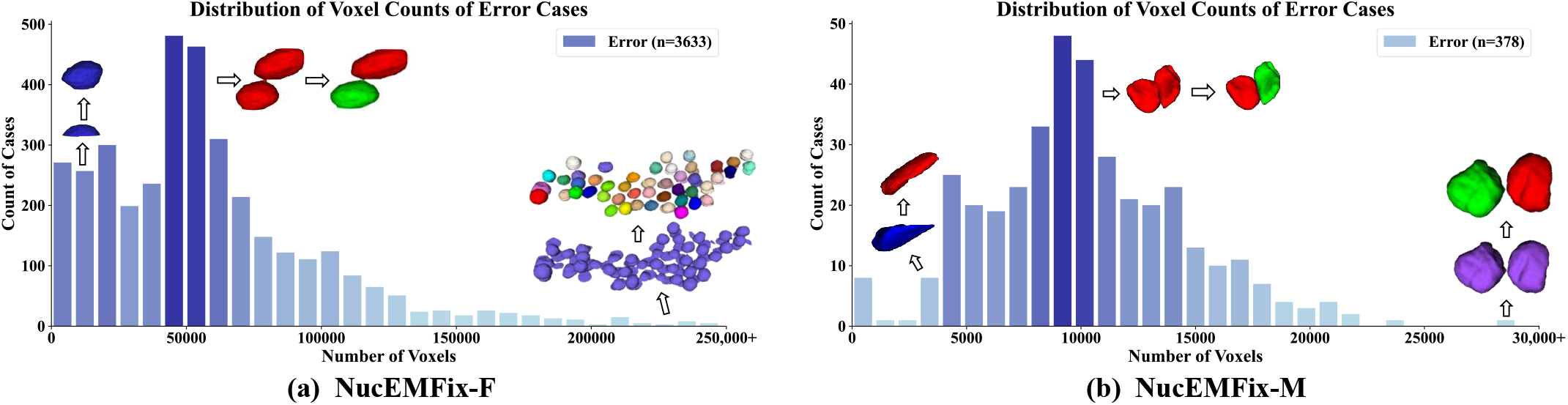
Long-tailed distribution of error case sizes in NucEMFix. Cases span the whole brain and vary widely in shape and size across three types: false merge, false split, and false merge&split.

## V. Experiments

### A. Experiment Settings

#### Evaluation Metrics

To evaluate performance, we adapt standard 3D instance segmentation metrics, precision, recall, and F1-score. A predicted nucleus and a ground-truth instance are considered a match if their Intersection-over-Union (IoU) is at least *τ*. Unless otherwise noted, we use *τ* = 0.75 [19].

#### Methods in Comparison

We compare our approach against four representative nucleus instance segmentation methods: *U-Net* [17], *nnU-Net* [18], *StarDist* [19], and *Cellpose* [20].

#### Implementation Details

All training data are derived from the FAFB and MICrONS datasets. We first exclude the NucEMFix error cases, then randomly select 5,000 clean nuclei as a base set for synthetic data generation. For training the reasoning module, these clean masks are used to synthesize 5,000 topological error cases as described in Sec. III-C. For shape refinement, 5,000 local 3D patches are extracted by cropping cubic volumes centered at each nucleus centroid (128^3^ for FAFB, 64^3^ for MICrONS) and stochastically degraded to generate corrupted-input/clean-target pairs. For baseline methods, all original nuclei masks within these patches are retained as training targets. All synthetic data are split into training and validation sets with an 8:2 ratio. All models are trained on a single NVIDIA A800 GPU using the Adam optimizer with a learning rate of 1*×*10^*−*4^, a batch size of 32, and up to 100 epochs with early stopping to prevent overfitting.

### B. Benchmark Results

#### Quantitative Results

Our evaluation covers two complementary comparison paradigms. Table II and Fig. 7 compares NuGraph against re-segmentation baselines (U-Net, nnU-Net, StarDist, Cellpose), which segment entirely from scratch without access to any prior mask information. Table IV further provides correction-specific baselines including pairwise matching and graph agglomeration methods, which is discussed in detail in Sec. V-C. NuGraph significantly outperforms all re-segmentation baselines across both subsets. On NucEMFix-F, NuGraph achieves an F1 of 87.99% (+8.58% over nnU-Net), with gains driven primarily by resolving false splits caused by axial gaps and false merges in dense clusters where ambiguous boundaries confound segmentation methods. On NucEMFix-M, most false-split and hybrid errors originate from severe misalignment where realignment preprocessing fails to restore structural continuity. Performance on these cases remains degraded across all methods, as instance reconstruction becomes infeasible without image evidence. Despite this, NuGraph achieves the highest overall F1 of 86.20%.

**TABLE I.** NucEMFix statistics. “Initial” counts released instances before curation. “Error-affected” counts nuclei impacted by each error type: FM (false merge), FS (false split), FM&S (false merge&split). Volume size in kilo-voxels.

| Subset | Volume (kvox) | Initial | Error-affected nuclei |  |  |  |
| --- | --- | --- | --- | --- | --- | --- |
|  |  |  | FM | FS | FM&S | Total |
| NucEMFix-F | $3.6 \times 8.2 \times 14.3$ | 106,978 | 5,258 | 1,031 | 1,012 | 7,301 |
| NucEMFix-M | $1.6 \times 4.1 \times 6.1$ | 77,475 | 711 | 20 | 26 | 757 |

**TABLE II.** Benchmark results on NucEMFix. Re-segmentation baselines (U-Net, nnU-Net, StarDist, Cellpose) segment from scratch; NuGraph reasons over existing primitives. Initial segmentation results show the severity of uncorrected errors.

| Method | NucEMFix-F |  |  | NucEMFix-M |  |  |
| --- | --- | --- | --- | --- | --- | --- |
|  | Pre (%) | Rec (%) | F1 (%) | Pre (%) | Rec (%) | F1 (%) |
| Initial Seg | 12.83 | 9.50 | 10.92 | 8.74 | 5.04 | 6.39 |
| U-Net [17] | 68.30 | 87.69 | 76.79 | 73.68 | 88.49 | 80.41 |
| nnU-Net [18] | 71.95 | 88.60 | 79.41 | 41.49 | 42.86 | 42.16 |
| StarDist [19] | 69.03 | 77.75 | 73.13 | 19.10 | 20.13 | 19.60 |
| Cellpose [20] | 36.34 | 73.24 | 48.58 | 44.72 | 56.61 | 49.97 |
| **Ours** | **86.50** | **89.53** | **87.99** | **82.45** | **90.32** | **86.20** |

**TABLE III.** Stage-wise ablation of NuGraph. S1: shape decomposition, S2: graph-based reasoning, S3: shape refinement. S1+S3 shows that shape refinement alone cannot substitute for relational reasoning.

| Stage |  |  | False Merge |  |  | False Split |  |  | False Merge&Split |  |  | All |  |  |
| --- | --- | --- | --- | --- | --- | --- | --- | --- | --- | --- | --- | --- | --- | --- |
| S1 | S2 | S3 | Pre (%) | Rec (%) | F1 (%) | Pre (%) | Rec (%) | F1 (%) | Pre (%) | Rec (%) | F1 (%) | Pre (%) | Rec (%) | F1 (%) |
| — | — | — | 6.87 | 3.37 | 4.52 | 19.91 | 41.59 | 26.92 | 13.24 | 8.89 | 10.64 | 12.83 | 9.50 | 10.92 |
| ✓ | — | — | 52.77 | 82.27 | 64.30 | 7.69 | 25.15 | 11.78 | 10.86 | 29.84 | 15.93 | 34.10 | 66.98 | 45.19 |
| ✓ | ✓ | — | 80.35 | 90.25 | 85.01 | 48.27 | 67.52 | 56.30 | 35.18 | 55.73 | 43.14 | 67.14 | 82.29 | 73.94 |
| ✓ | — | ✓ | 86.58 | 91.15 | 88.80 | 64.85 | 75.83 | 69.91 | 57.60 | 65.12 | 61.13 | 79.07 | 85.38 | 82.10 |
| ✓ | ✓ | ✓ | **92.29** | **94.19** | **93.23** | **75.48** | **85.39** | **80.13** | **68.52** | **69.47** | **68.99** | **86.50** | **89.53** | **87.99** |

**TABLE IV.** Reasoning architecture ablation. Local pair-matching methods vs. our global graph-based reasoning approach. *∗* denotes the multicut-based version of EdgeNetwork.

| Method | False Merge |  |  | False Split |  |  | False Merge&Split |  |  | All |  |  |
| --- | --- | --- | --- | --- | --- | --- | --- | --- | --- | --- | --- | --- |
|  | Pre (%) | Rec (%) | F1 (%) | Pre (%) | Rec (%) | F1 (%) | Pre (%) | Rec (%) | F1 (%) | Pre (%) | Rec (%) | F1 (%) |
| *Pair-Matching* |  |  |  |  |  |  |  |  |  |  |  |  |
| PointNet++ [15] | 84.76 | 78.35 | 81.43 | **76.05** | **85.88** | **80.67** | 55.81 | 43.68 | 49.00 | 79.91 | 74.58 | 77.15 |
| EdgeNetwork [16] | 89.22 | 92.65 | 90.90 | 67.49 | 78.68 | 72.65 | 61.14 | 67.00 | 63.93 | 81.88 | 87.14 | 84.43 |
| *Graph Reasoning* |  |  |  |  |  |  |  |  |  |  |  |  |
| EdgeNetwork* [16] | 89.41 | 92.81 | 91.08 | 67.57 | 78.77 | 72.74 | 61.74 | 67.60 | 64.54 | 82.36 | 87.58 | 84.88 |
| Ours w/o $\mathcal{L}_{\text{contrast}}$ | 90.92 | 91.76 | 91.34 | 74.16 | 84.70 | 79.08 | 61.69 | 67.00 | 64.23 | 84.94 | 86.46 | 85.67 |
| **Ours** | **92.29** | **94.19** | **93.23** | 75.48 | 85.39 | 80.13 | **68.52** | **69.47** | **68.99** | **86.50** | **89.53** | **87.99** |

**TABLE V.**
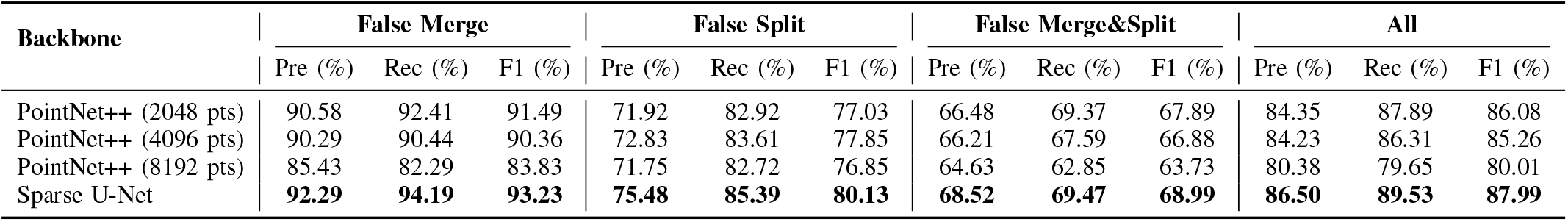
3D backbone comparison for the reasoning module. PointNet++ with fixed point sampling vs. Sparse U-Net.

**Fig. 7.**
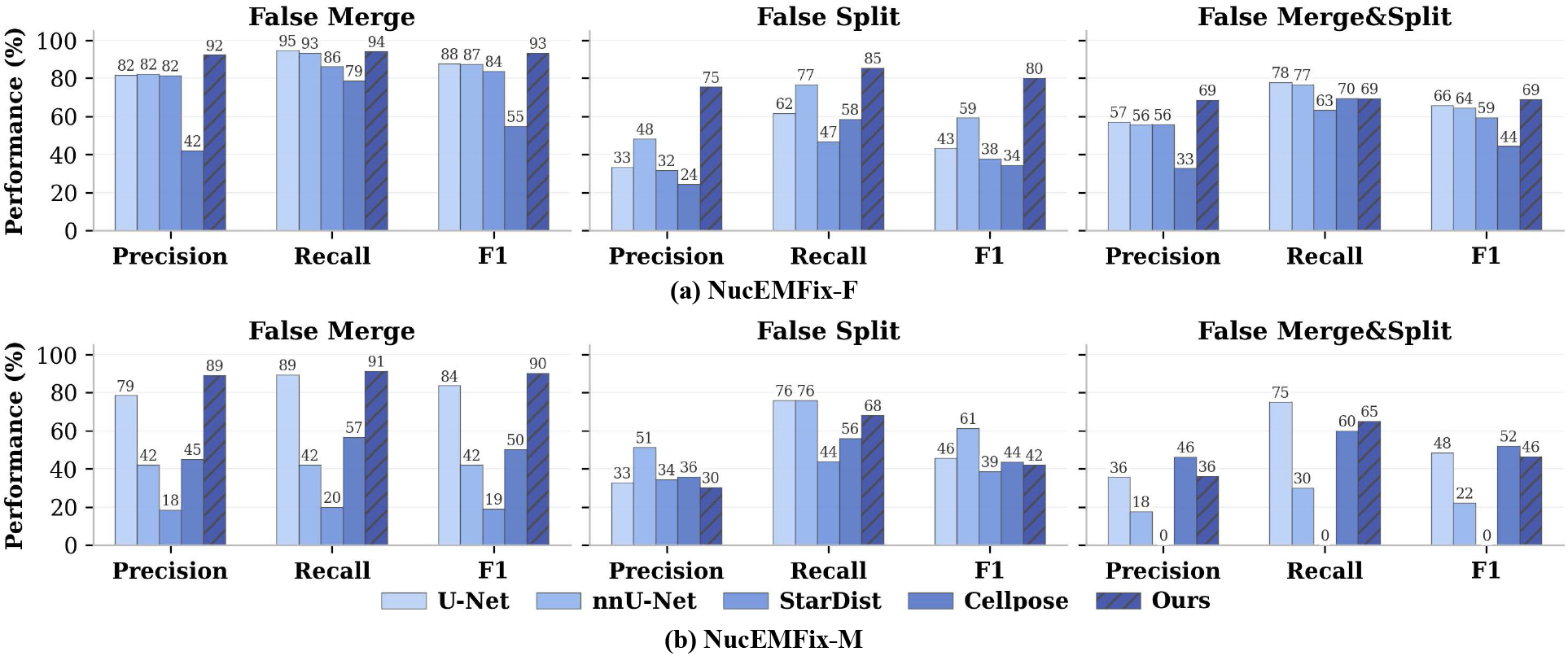
Per-error-type F1 on NucEMFix across all methods, broken down by false merge, false split, and merge&split.

#### Qualitative Results

Fig. 8 shows representative correction results. U-Net produces non-biological sheet-like false splits. nnU-Net fails to correct false merges and yields eroded segmentation. StarDist cannot adapt to irregular boundaries in NucEMFix due to its star-convexity constraint. Cellpose produces fragmented results, as its gradient flows are disrupted by misalignment and noise. All baselines struggle with false splits from missing slices, as they cannot reason across vacant axial layers. In contrast, NuGraph achieves accurate correction across all error types by reasoning globally over primitives and restoring complete morphology.

**Fig. 8.**
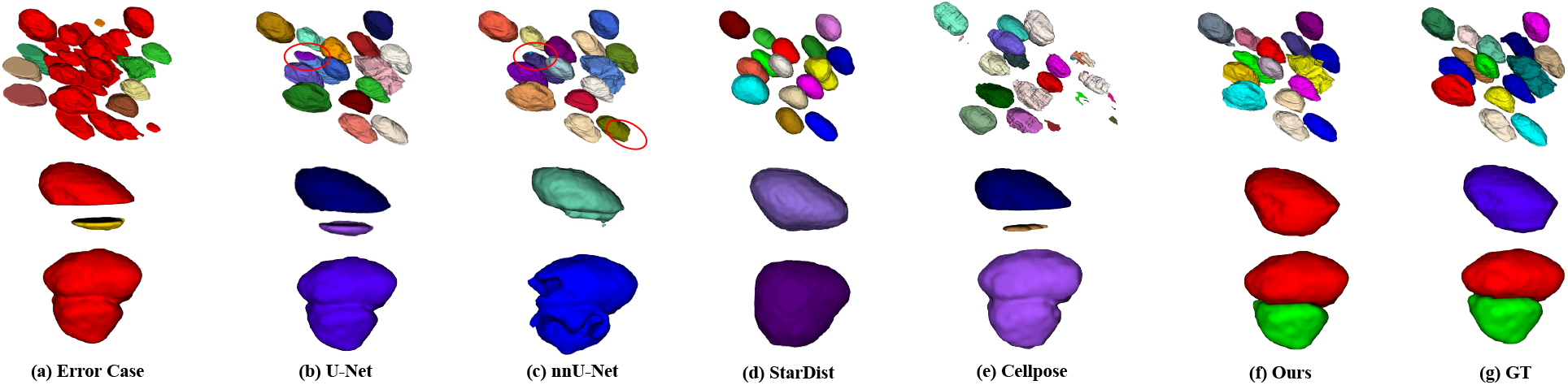
Qualitative correction results on NucEMFix. Different colors indicate different instances. Top: false merge&split with misalignment; middle: false split from missing axial slices; bottom: false merge.

### C. Ablation Studies

#### Stage-Wise Integration

As shown in Tab. III, we progressively integrate each stage. The initial segmentation yields F1 of 10.92%. Shape decomposition (S1) boosts F1 to 45.19% by decomposing dense clusters. Adding graph-based reasoning (S2) brings a 28.75% leap to 73.94%, establishing correct topology. The full NuGraph reaches 87.99%. The S1+S3 variant (skipping reasoning) achieves 82.10%—higher than S1+S2 alone on geometrically simple cases, but falling short of the full model on complex merge–split scenarios where shape refinement on raw primitives lacks topologically correct grouping. S2 and S3 are complementary: reasoning fixes topology and refinement smooths morphology.

#### Pair-Matching vs. Graph Reasoning

We compare local pair-matching baselines—PointNet++ [15] and EdgeNetwork [16], both widely used for neuron proofreading—against our graph-based approach (Tab. IV). Since these baselines were originally designed for neuron fragments, we re-implemented them and re-trained from scratch on the NucEMFix synthetic data to ensure they are adapted to the nuclei domain. For a controlled comparison, all methods share the same primitive decomposition, candidate-edge threshold, training data, and SDF refinement; only the reasoning module differs. Pair-matching methods predict pairwise connectivity and group primitives via union–find on the predicted links. EdgeNetwork^∗^ replaces union–find with multicut optimization [31] for a globally consistent partitioning, but its edge weights are still derived from pairwise features alone, limiting reasoning over densely entangled regions. PointNet++ performs reasonably on isolated false splits but struggles in dense merge–split hybrids (F1: 49.00%), where a primitive’s identity cannot be resolved without broader context. EdgeNetwork improves to 84.43% via learned edge features, and its multicut variant adds only a marginal gain (84.88%), confirming that the bottleneck lies in pairwise feature expressiveness rather than the grouping strategy. Graph attention reasoning raises F1 to 85.67% by aggregating neighborhood context across the entire cluster, and contrastive alignment adds a further 2.32% (87.99%) by aligning primitive features with proposal-level semantics.

#### Robustness to Case Complexity

Fig. 9 reports F1 as a function of the number of primitives per error case on the False Merge&Split subset of NucEMFix-F. For simple cases with 2–5 primitives, all methods perform comparably, as pairwise decisions suffice when the grouping structure is uncomplicated. However, performance diverges sharply as complexity grows: PointNet++ drops from 78.3% at 5 primitives to 42.3% at 11+, and EdgeNetwork from 82.2% to 59.0%, reflecting their inability to resolve ambiguous groupings without global context. NuGraph degrades more gracefully, maintaining 65.5% F1 even at 11+ primitives, as graph attention aggregates neighborhood evidence across all primitives rather than reasoning over isolated pairs.

**Fig. 9.**
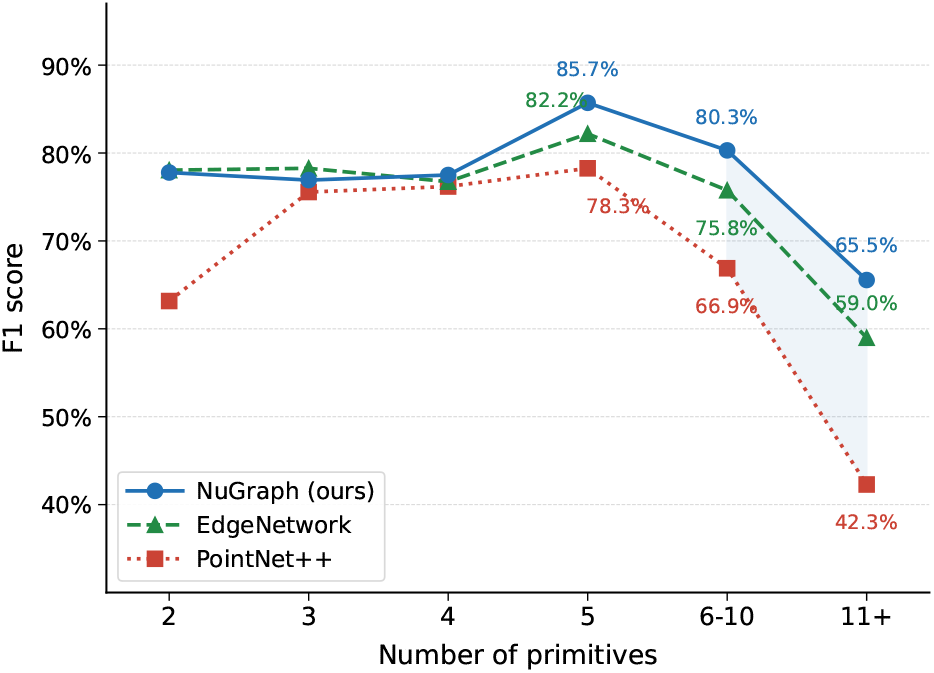
F1 vs. primitive count per case on the False Merge&Split subset of NucEMFix-F. NuGraph degrades more gracefully as complexity grows.

#### Effect of Contrastive Alignment

Fig. 10 visualizes primitive feature embeddings: without contrastive learning, similarity patterns are diffuse; with contrastive alignment, intra-group similarity becomes compact and inter-group separation clear, providing more reliable edge cues for graph reasoning.

**Fig. 10.**
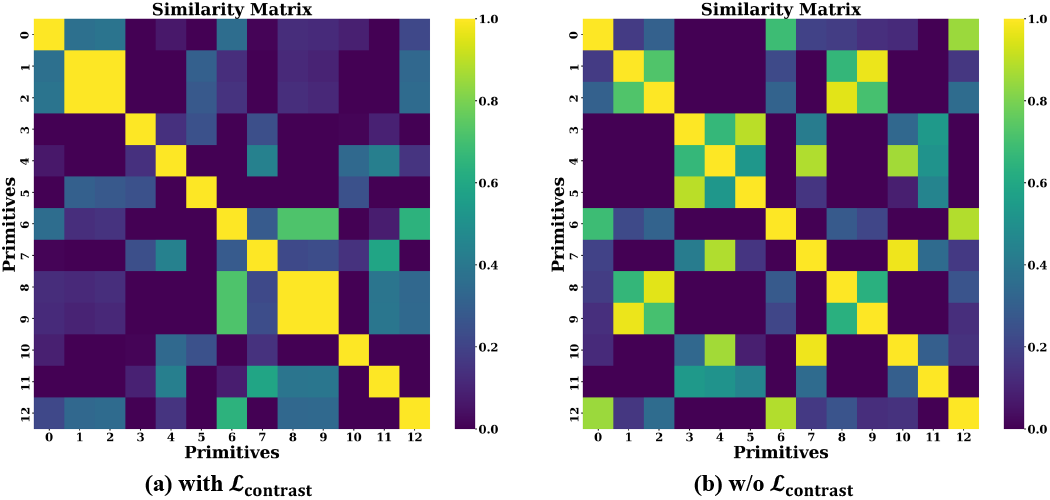
Pairwise similarity of primitive embeddings, with and without primitive–proposal contrastive learning. Ground-truth grouping: {1},{2,3},{4},{5},{6},{7},{8,9},{10},{11},{12}.

#### 3D Backbone Choice

Tab. V compares 3D backbones for the reasoning module. PointNet++ with fixed-size sampling shows limited robustness under long-tailed error distributions, as small cases are over-represented and larger regions suffer from redundant padding. Sparse U-Net processes variable-length surfaces without uniform sampling and benefits from a larger receptive field, enabling more effective global reasoning.

#### Shape Refinement

Table VI compares mask-only reconstruction against SDF-guided completion. SDF supervision improves F1 on false splits from 78.61% to 80.13% (+1.52%) and overall F1 by +1.35%, confirming that continuous SDF representation helps recover fragmented nuclei with missing slices.

**TABLE VI.** Shape completion ablation. Mask-only reconstruction vs. SDF-guided completion.

| Method | False Merge |  |  | False Split |  |  | False Merge&Split |  |  | All |  |  |
| --- | --- | --- | --- | --- | --- | --- | --- | --- | --- | --- | --- | --- |
|  | Pre (%) | Rec (%) | F1 (%) | Pre (%) | Rec (%) | F1 (%) | Pre (%) | Rec (%) | F1 (%) | Pre (%) | Rec (%) | F1 (%) |
| w/o SDF (Mask-only) | 91.40 | 93.95 | 92.66 | 73.33 | 84.70 | 78.61 | 67.11 | 70.36 | 68.69 | 84.76 | 88.60 | 86.64 |
| Ours (with SDF) | **92.29** | **94.19** | **93.23** | **75.48** | **85.39** | **80.13** | **68.52** | **69.47** | **68.99** | **86.50** | **89.53** | **87.99** |

#### Hyperparameter Sensitivity

We evaluate sensitivity to graph construction threshold *d*, contrastive weight *λ*_contrast_, and SDF weight *λ*_sdf_ (Fig. 11). Performance is stable across a wide range for all three. Best results are at *d*=7, *λ*_contrast_=1, *λ*_sdf_ =1.

**Fig. 11.**
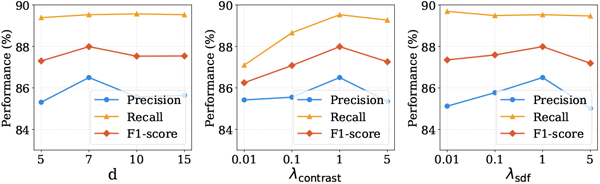
Sensitivity to graph construction threshold ***d***, contrastive weight ***λ***_contrast_, and SDF weight ***λ***_sdf_. F1 is stable across a broad range.

### D. More Analysis

#### Stability on Error-Free Nuclei

To verify that NuGraph does not over-correct accurate segmentations, we evaluated on 30,000 (FAFB) and 10,000 (MICrONS) correctly annotated nuclei. Only one nucleus was erroneously split in FAFB and four underwent minor modifications in MICrONS, confirming that NuGraph preserves segmentation quality brain-wide.

#### Inference Cost

Table VII reports the inference time of NuGraph on NucEMFix-F, broken down by error type and pipeline stage. Across all 3,633 error cases, NuGraph completes correction in approximately 85.2 minutes on a single NVIDIA A800 GPU running single-threaded sequential inference. The shape decomposition stage (S1) dominates runtime due to EDT computation and slice realignment, while graph reasoning (S2) and SDF refinement (S3) contribute smaller overhead. In contrast, manual curation of the same cases required over 100 annotator-hours, yielding a speedup of more than 100*×* relative to human correction.

**TABLE VII.**
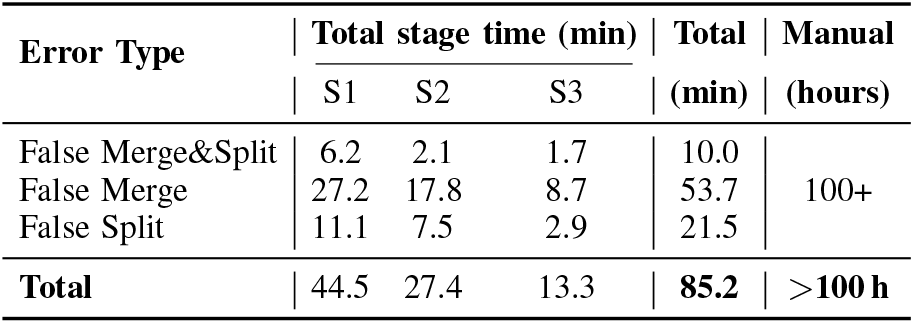
Inference time of NuGraph on NucEMFix-f per error type. S1: shape decomposition and alignment; S2: graph-based reasoning; S3: sdf shape refinement.

| Error Type | Total stage time (min) |  |  | Total<br>(min) | Manual<br>(hours) |
| --- | --- | --- | --- | --- | --- |
|  | S1 | S2 | S3 |  |  |
| False Merge&Split | 6.2 | 2.1 | 1.7 | 10.0 | 100+ |
| False Merge | 27.2 | 17.8 | 8.7 | 53.7 |  |
| False Split | 11.1 | 7.5 | 2.9 | 21.5 |  |
| **Total** | **44.5** | **27.4** | **13.3** | **85.2** | **>100 h** |

#### Failure Cases

Fig. 12 shows failure cases. (a) A nucleus with a deep invagination: the reasoning module fails to merge the two primitives, as the irregular shape lacks the geometric cues needed for confident grouping. (b) Two nuclei with an extremely large contact surface: the module incorrectly predicts high connectivity, producing a false merge. (c) Severe misalignment with illumination artifacts: preprocessing fails to restore continuity, leaving the reasoning module without usable structural input. Such cases are prevalent in NucEMFix-M.

**Fig. 12.**
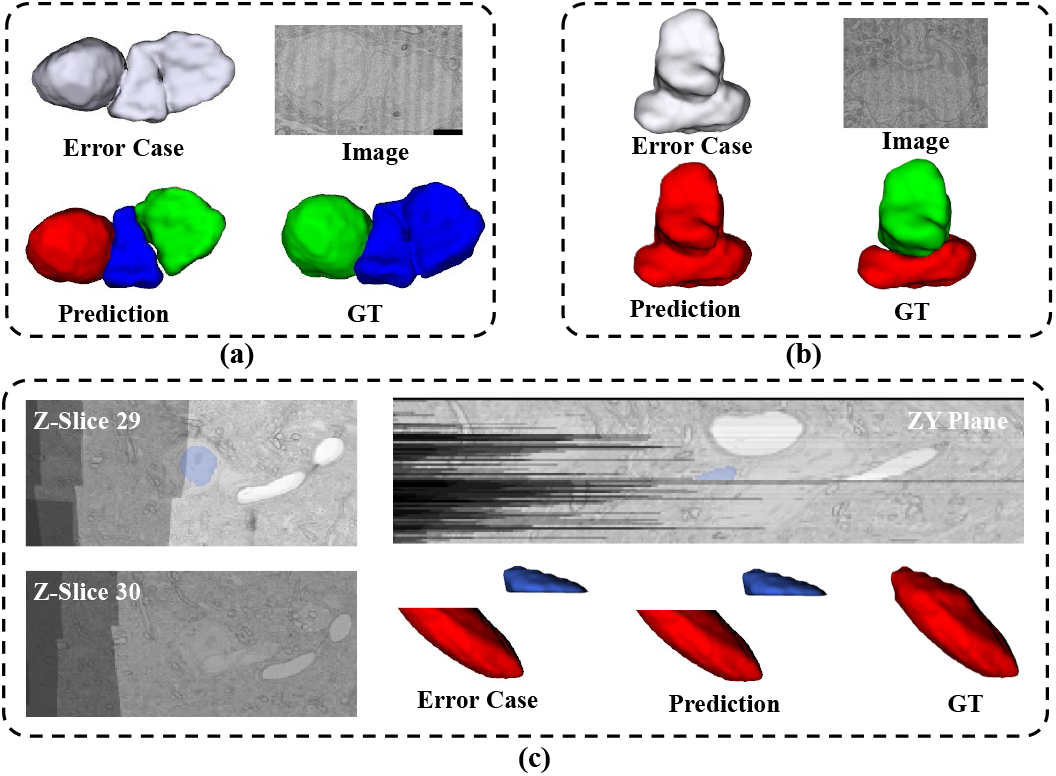
Failure cases. (a) Deep invagination misread as two primitives; (b) large contact surface causing a false merge; (c) severe misalignment with illumination artifacts breaking structural continuity.

## VI. Discussion

### Why dense clusters remain the hardest regime

Across both subsets, residual errors concentrate on tightly packed, irregular nuclei—e.g., perivascular populations in MICrONS and large merge clusters in FAFB—where local boundary cues are weak and primitive decomposition is most ambiguous. Our graph reasoning mitigates but does not eliminate this regime: Fig. 9 shows graceful degradation rather than a hard ceiling, suggesting better primitive proposals—not richer reasoning— are the next bottleneck.

### Cross-dataset gap

The performance asymmetry between NucEMFix-F and NucEMFix-M stems mainly from acquisition-level differences. MICrONS has a larger fraction of cases dominated by severe misalignment and illumination artifacts that destroy structural continuity, leaving the geometry-only reasoning module without usable input (cf. Sec. V-B and the failure cases in Fig. 12). This points to image-conditioned reasoning as a clear next step.

### Implications for downstream analysis

Beyond raw F1, the SDF refinement head restores smooth, biologically plausible morphology—a prerequisite for downstream cell typing and morphological analysis. Combined with the brain-wide stability evaluation (Sec. V-D), this suggests NuGraph can be deployed across entire connectomic volumes without quality regression.

### Limitations and Future Work

The data engine synthesizes errors from clean masks alone, restricting reasoning to geometric cues; severe imaging artifacts and missing slices that destroy structural continuity remain hard cases. Future work will couple error generation to raw EM imagery and explore image-conditioned graph reasoning at no added inference cost.

## VII. Conclusion

We presented NuGraph, a graph-based framework that corrects 3D nuclei segmentation by decomposing errors into atomic primitives and reasoning over them with graph attention and primitive–proposal contrastive alignment. This formulation captures the global topology of dense clusters that local pairwise methods miss, while an SDF refinement head restores smooth morphology. Trained entirely on self-supervised synthetic errors, NuGraph eliminates the need for manual error annotation. We also released **N****uc****EMFix**, the first brain-wide EM benchmark for nuclei correction with 8,000+ error cases across FAFB and MICrONS. NuGraph attains 87.99% F1 on NucEMFix-F and 86.20% on NucEMFix-M, outperforming all baselines and cutting curation effort by over*×*100 relative to manual proofreading. By releasing both the framework and a standardized correction benchmark, we hope NucEMFix will serve the broader connectomics community as a common testbed for measuring future last-mile correction methods at brain scale.

